# Water Nanoconfined in a Hydrophobic Pore: MD Simulations and Water Models

**DOI:** 10.1101/2021.07.28.453939

**Authors:** Charlotte I. Lynch, Gianni Klesse, Shanlin Rao, Stephen J. Tucker, Mark S. P. Sansom

## Abstract

Water molecules within biological ion channels are in a nano-confined environment and therefore exhibit novel behaviours which differ from that of bulk water. Here, we investigate the phenomenon of hydrophobic gating, the process by which a nanopore may spontaneously de-wet to form a ‘vapour lock’ if the pore is sufficiently hydrophobic and/or narrow. Notably, this occurs without steric occlusion of the pore. Using molecular dynamics simulations with both additive and polarisable (AMOEBA) force fields, we investigate this wetting/de-wetting behaviour in the TMEM175 ion channel. We examine how a range of rigid fixed-charge (i.e. additive) and polarisable water models affect wetting/de-wetting in both the wild-type structure and in mutants chosen to cover a range of nanopore radii and pore-lining hydrophobicities. Crucially, we find that the rigid fixed-charge water models lead to similar wetting/de-wetting behaviours, but that the polarisable water model resulted in an increased wettability of the hydrophobic gating region of the pore. This has significant implications for molecular simulations of nano-confined water, as it implies that polarisability may need to be included if we are to gain detailed mechanistic insights into wetting/de-wetting processes. These findings are of importance for the design of functionalised biomimetic nanopores (for e.g. sensing or desalination), as well as for furthering our understanding of the mechanistic processes underlying biological ion channel function.

## Introduction

The transport behaviour of water and other liquids in nanoscale pores is of both fundamental and technological importance ^1^. Nanoscale pores across membranes are of especial interest ^2^. In biological systems, channels are protein nanopores which enable the flow of water molecules and/or ions across cell membranes ^3^. The behaviour of water molecules within nanoscale pores differs in several respects from that of bulk water ^4–7^. In particular, ion channel proteins and nanopores may possess a hydrophobic gate (Fig. 1A). This is a constricted region of the pore which is lined with hydrophobic residues in which de-wetting may occur to form a “vapour lock”^8–9^ which in turn presents an energetic barrier to ion permeation and thereby functionally closes the channel. Therefore, in these ion channels, the hydrophobic gate closes the pore to the passage of water and ions ^10^ without the need for steric occlusion of the pore. The presence of hydrophobic gates in several ion channel species has been inferred from both computational and experimental studies. Hydrophobic gating has also been designed into synthetic nanopores e.g. ^11^. Thus, a detailed mechanistic understanding of hydrophobic gating is relevant both to studies of ion channel structure/function relationships ^12^, and also to the design of gating functionality ^13^ into synthetic nanopores ^14–15^.

**Figure 1:**
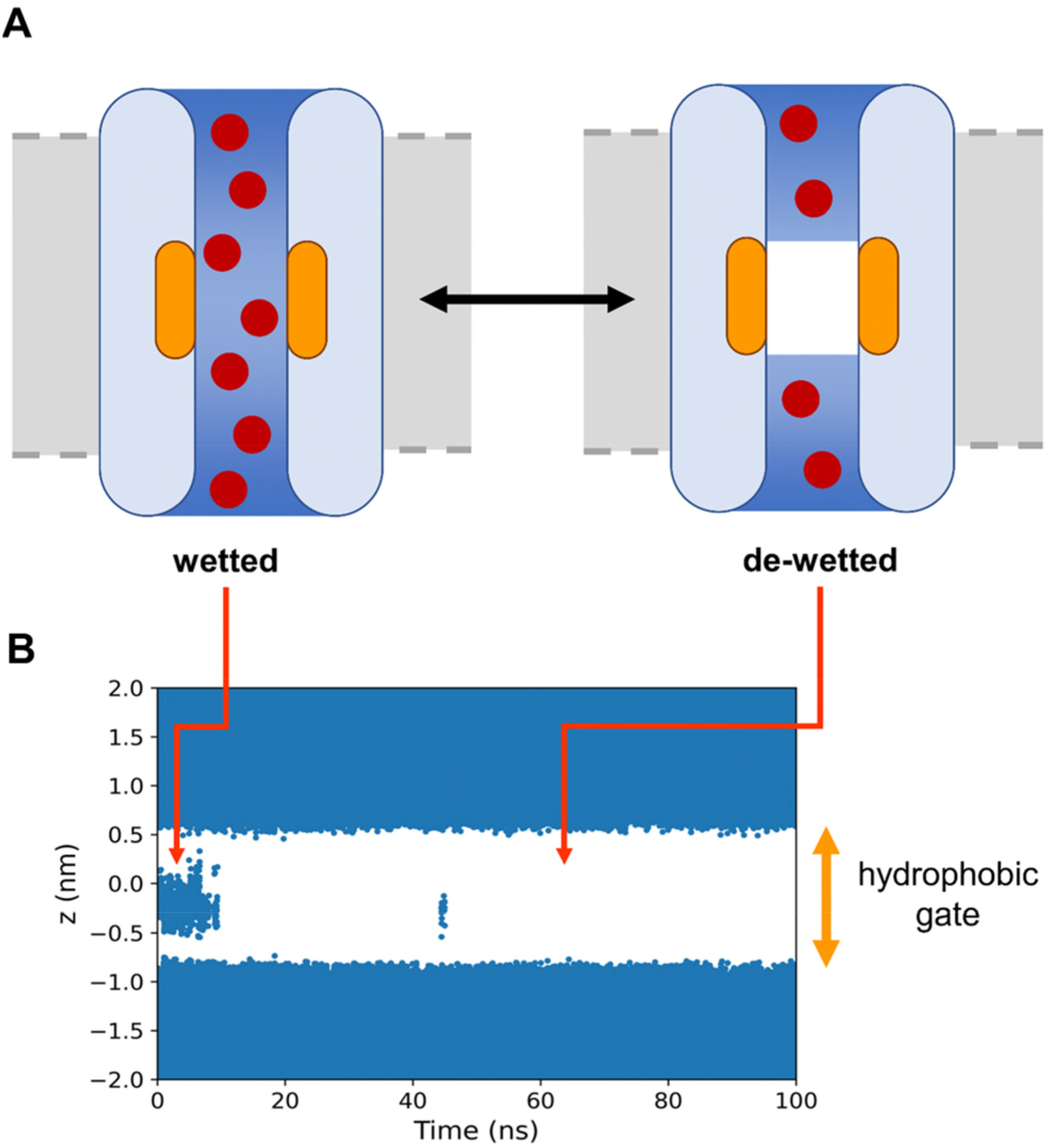
**A** The principle of hydrophobic gating: in the wetted state, both water (medium blue) and ions (red) can pass through the ion channel (pale blue) and the hydrophobic gate (orange); whereas in the de-wetted state, the channel, though not physically occluded, is closed to the passage of ions. **B** An example of simulated de-wetting of a hydrophobic gate. The trajectories of water molecules (dark blue) around and initially located in a hydrophobic gating region (indicated by the orange arrow, centred at *z* = 0) are shown projected onto the *z* axis. The data are taken from a simulation of the TMEM175 channel protein using the TIP4P water model (see main text for details; also see SI Fig. S1).

The behaviour of a hydrophobic gate has been shown to depend critically on the radius of the pore and on the hydrophobicity of the pore lining ^16–19^. Furthermore, in addition to these two structural parameters, the application of an electric field ^20–21^ or pressure ^22–23^ can also cause the pore to wet, thereby opening an otherwise closed channel to the passage of water and ions.

Molecular dynamics (MD) simulations have been widely applied to explore the parameters affecting hydrophobic gating and the wetting/de-wetting behaviour both of ion channels and of simplified models of nanopores (see e.g. ^6,12^ for two recent reviews). By measuring the number density of water molecules within the gating region, an associated energetic barrier to wetting can be estimated ^24^. This has been used to characterise the likely functional state of nearly 200 experimentally determined ion channel structures ^19^, leading to a more global analysis of the dependence of hydrophobic gating behaviour (in terms of the energetic barrier to wetting) on channel structure. MD simulations have also been used to explore the effect of pore lining mutations on hydrophobic gating in e.g. the bestrophin-1 chloride channel ^25^.

When conducting MD simulations of water behaviour in nanopores and channels, a crucial but not yet widely investigated consideration is the choice of water model. For example, recent simulations have examined the effect of varying water models on the behaviour of water and ions within cyclic peptide nanopores ^26^ and within ligand gated ion channels ^27^.

Due to the competing effects of hydrogen bonding, van der Waals and other intermolecular interactions, water is a particularly difficult molecule to model accurately. To date, there are over 130 different water models for use in MD simulations and related calculations, each with a range of physical properties which are comparable to experimental values ^6^. Typically, these water models are developed to accurately reproduce one or more experimental properties of bulk water. In the case of water in hydrophobic gates, however, water molecules are confined within a region of ~1 nm diameter and ~2 nm in length. This means that there are likely to be only a small number of water molecules in the hydrophobic gating region, with a maximum of ~50 for a cylinder of this volume. Water confined within such a nanoscale channel is unlikely to behave as it does in bulk, e.g. in terms of its dielectric behaviour ^7^. Consequently, it is not certain that water models developed to reproduce bulk behaviour will be directly transferable to such nanoconfined environments. Initial studies on the pore domain of the 5-HT3 receptor suggest that the choice of water model can have an effect on the wetting/de-wetting behaviour, particularly in the case of structures which are in an intermediate open/closed state ^27^. In particular, there are indications that the use of more sophisticated water models such as polarisable water models (the dipole moment of which will be responsive to changes in local environment) may be required to more adequately capture the interactions within hydrophobic gates. Although polarisable models are more computationally expensive, the development of more powerful computing facilities and of improved software enables the use of such models for complex biological systems ^28–29^. Thus, recent studies have explored the influence of polarisable models on e.g. simulations of the interactions of ions with lipid bilayer membranes ^30^ and with simple model ion channels e.g. ^31^. Various rigid fixed-charge water models have also been assessed for ligand binding in host/guest systems ^32^ and for water behaviour in peptide nanotubes ^26^. Wetting/de-wetting of a hydrophobic gate provides an extreme case of non-bulk behaviour of water when nanoconfined in a pore. We have therefore used a well-defined hydrophobic gate, present in the TMEM175 ion channel ^25, 33–36^, to explore this for different rigid fixed-charge water models in comparison with the AMOEBA ^37–38^ polarisable water model.

In the current study we use the prokaryotic CmTMEM175 structure ^33^ to provide a model of a highly nanoconfined environment within the hydrophobic gate of an ion channel. We investigate the effects of changing both the pore-lining residues and the water model employed in the simulations on wetting/de-wetting behaviour of this hydrophobic gate. This region of the channel is formed by three hydrophobic rings (each with four-fold symmetry): one ring of isoleucine residues (residue Ile23) followed by two of leucine residues (Leu27 and Leu30 respectively). We explore how mutating this hydrophobic gate to either three rings of valines (slightly smaller hydrophobic sidechains), or alanines (hydrophobic but much smaller sidechains), or asparagines (hydrophilic sidechains) affects the water behaviour. We also investigate the transferability of different rigid fixed-charge (additive) and polarisable water models for simulating water behaviour within the hydrophobic gate, i.e. we evaluate whether the different water models produce consistent wetting/de-wetting behaviours. Our results provide mechanistic insights into the behaviour of water within a highly nanoconfined pore. This is of direct relevance to the ongoing interest in nanoconfined water as studied either experimentally (e.g. ^39–40^) or computationally (e.g. ^7,41^).

## Results and Discussion

### Channel System and Simulations

In this study we used the prokaryotic CmTMEM175 (PBD: 5VRE; 3.3 Å resolution) ^33^ as a model for a pore with a hydrophobic gate which can be wetted/de-wetted ^25^. In this channel structure the first transmembrane helix from each of four subunits line a central pore with a constriction (with a minimum radius of < 0.15 nm on the crystal structure) formed by three rings of hydrophobic sidechains: Ile23, Leu27 and Leu30 (see Fig. 2). In two other TMEM175 channels these rings of residues are either conserved (in hTMEM175, PDB: 6WC9) or replaced by three rings of leucines (in MtTMEM175, PDB: 6HD8). However, the structural conservation of this region is less clear-cut ^33–35^, as might be expected for a hydrophobic gating region.

**Figure 2:**
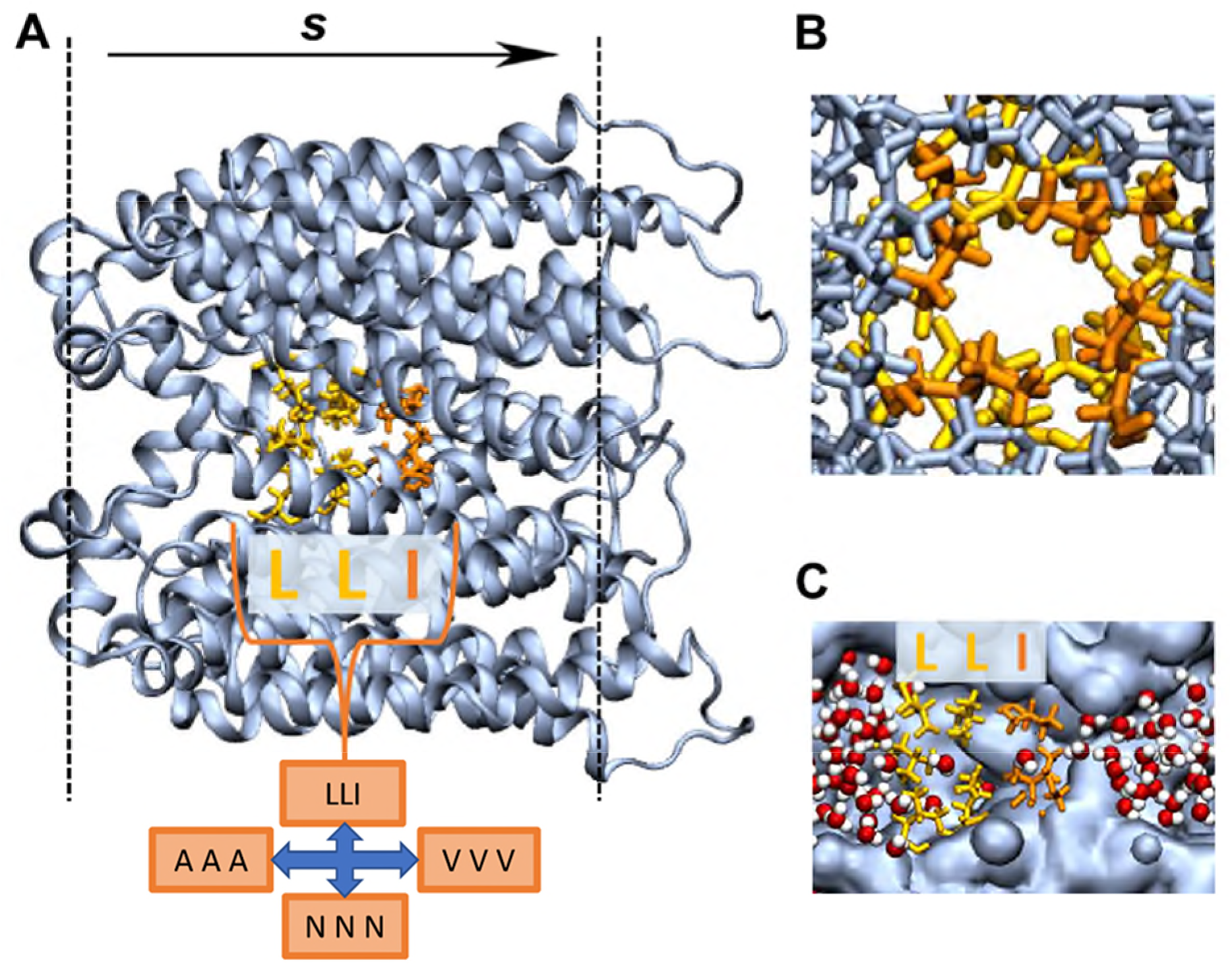
**A** The simulation set-up with the TMEM175 protein in blue and the wild-type hydrophobic gating residues highlighted in yellow (LEU) and orange (ILE). The LEU and ILE residues in the hydrophobic gate (LLI) are substituted for ALA, ASN and VAL respectively to create three mutants (AAA, NNN and VVV respectively). The position of the lipid headgroups in the bilayer are denoted by the dashed black line, and *s* indicates the path direction through the centre of the pore. **B** A close-up looking down the wild-type pore. **C** A slice through the pore with water (red spheres for oxygen, white spheres for hydrogen) exiting the hydrophobic gating region of the pore surface (blue).

In order to explore systematically some of the parameters affecting hydrophobic gating (e.g. pore radius and hydrophobicity), the CmTMEM175 structure (PDB code 5VRE) was embedded within a phosphatidylcholine (PC) bilayer and solvated with a ~0.15 M NaCl aqueous solution. In these simulations, backbone restraints were applied in order to maintain the experimentally observed polypeptide backbone conformation whilst allowing a degree of sidechain flexibility/mobility. In initial simulations of the wild type (i.e. native) protein the gating region was seen to be sufficiently narrow and hydrophobic that the pore spontaneously de-wetted early on in simulations (Fig. 1B and SI Fig. S1A).

As can be seen (Fig. 2), the hydrophobic gate in the CmTMEM175 structure is formed by a constricted region of three rings of hydrophobic sidechains, forming an ‘ILL’ motif. In order to explore the sensitivity of wetting/de-wetting behaviour to the nature of this motif, these residues were *in silico* mutated to either three rings of valines (VVV), i.e. hydrophobic sidechains but smaller than isoleucine or leucine, or three rings of alanines (still hydrophobic, but substantially smaller sidechains), or three asparagines (the sidechain of which is the same size as that of leucine, but is polar and able to form H-bonds) – see Fig. 2A. For both the wild type and the three mutant channels we conducted simulations with various water models (see below). Simulations were of 100 ns duration with 6 repeats for fixed-charge water models and of up to 50 ns duration with 3 repeats for the polarisable AMOEBA model.

### Effect of Mutations on Wetting/De-wetting

In simulations of the wild-type channel, the hydrophobic gate is associated with a constriction of radius down to 0.1-0.2 nm (Fig. 3A). This is comparable to the radius of a water molecule (0.14 nm). The sidechains of the isoleucines and leucines result in a highly hydrophobic lining. Thus, in simulations using the TIP4P/2005 water model and OPLS all-atom protein force field (see Methods below), the gating region de-wets within the first ~10 ns for most of the repeats (SI Fig. S1B) and remains relatively dry for the remainder of each of the simulations. However, in some repeats of the simulations, a water molecule is temporarily trapped within the gate, due to the tight constrictions of ~0.1 nm at each end of the gate. Consequently, the time averaged water density (Fig. 3B) does not completely fall to 0 nm^−3^ within the hydrophobic gate of the wild-type channel (this will be discussed in more detail below). The energetic barrier to water of this region (Fig. 3C) is 13.5 (± 4.8) k_B_T (averaged over the 6 repeats, ± standard deviation). This conformation of the pore therefore clearly corresponds to a functionally closed state.

**Figure 3:**
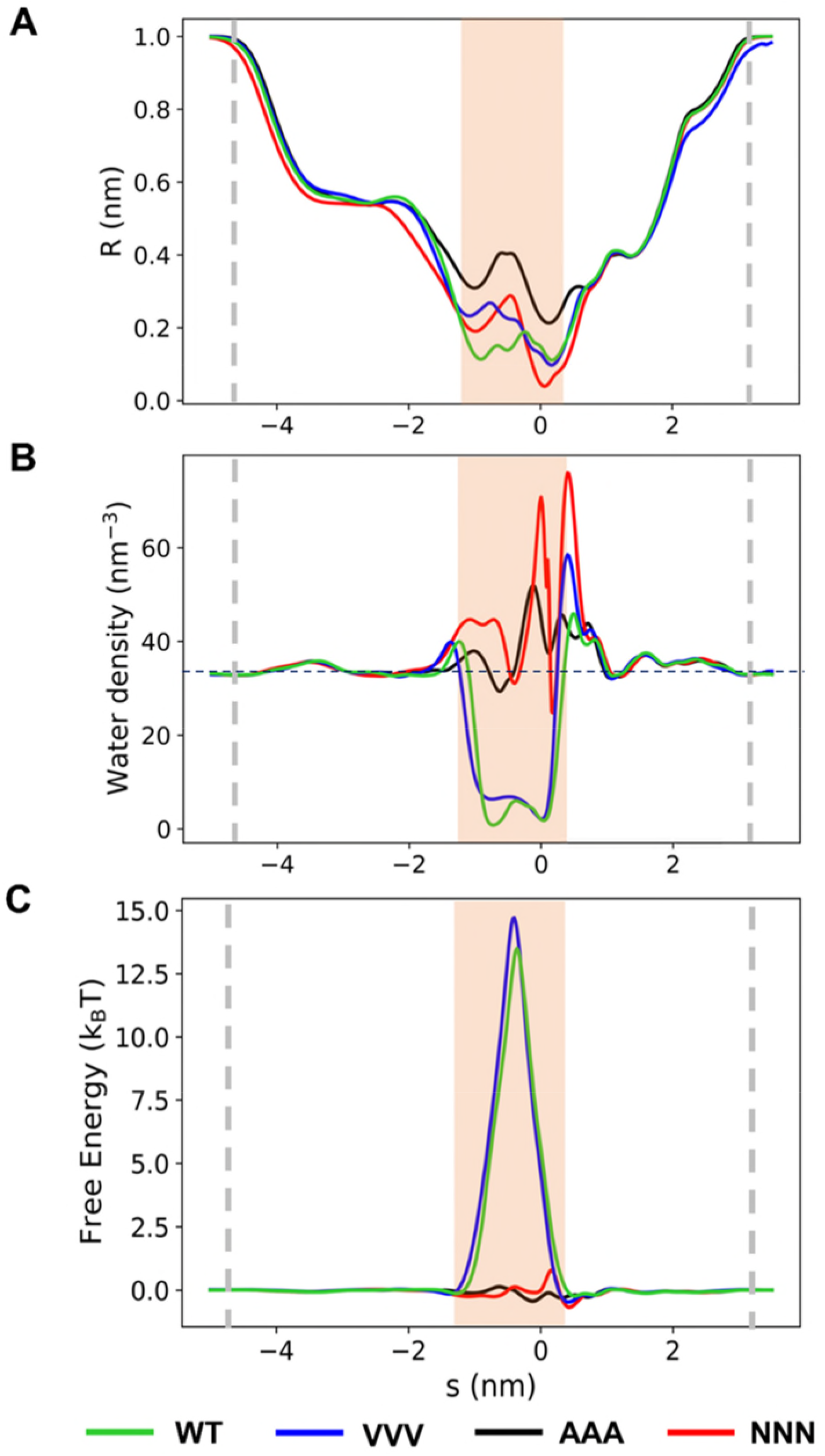
Profiles of pore radius (**A**), water density (**B**) and water free energy (**C**) as a function of distance along the pore centre line, s. The profiles for the wild-type structure (WT, green) and AAA (black), NNN (red) and VVV (blue) mutants are shown, each averaged over the last 90 ns of the simulation and subsequently averaged over 6 independent repeats. The water model TIP4P/2005 was used for all simulations. The hydrophobic gating region is indicated by the orange band, and the approximate positions of the lipid head groups are denoted by the grey dashed lines.

The constriction of the pore in the ‘VVV’ mutant remained hydrophobic but was a little wider (by ~0.1 nm at the lower/extracellular (i.e. *s* = −1.3 nm) end of the gate (Fig. 3A). Despite this small increase in radius (which was sufficient to prevent trapping of water molecules, with any water molecules present in the hydrophobic gate at the start of the simulation exiting via the wider leucine side) the pore remained de-wetted (Fig. 3B), with an energetic barrier comparable to that of the wildtype channel (Fig. 3C).

In contrast, although the AAA mutant preserved the hydrophobic nature of the pore lining, it was wider, with two minima (Fig. 3A) of ~0.2 and ~0.3 nm radius. This was sufficient for the pore to remain fully wetted (Fig. 3B), thus presenting no barrier to ion permeation (Fig. 3C). The more polar/hydrophilic mutant, NNN, had a pore radius profile comparable to that of the VVV mutant (Fig. 3A). However, the hydrophilic lining provided by the Asn (NNN) sidechains (to which water can form H-bonds) was such that the pore remained fully wetted (Fig. 3B) with no energetic barrier to water permeation (Fig. 3C).

### Effect of Additive Water Model on Wetting/De-wetting in the Wild-type Channel

The results discussed so far employed the additive (rigid fixed-charge) TIP4P/2005 water model, which has been shown to perform well over a range of (bulk) water properties ^42–43^ and has been used in recent studies of nanoconfined water e.g. ^44^. To explore the robustness of these results to variations in additive water models, we investigated the effect of five different models on the de-wetting behaviour in the hydrophobic gate of wild-type CmTMEM175 structure (Fig. 4). These models were chosen to cover a broad range of model types (see e.g. ^6,45^ for useful summaries). TIP3P and SPC/E are well-established 3-point water models with differing geometries: TIP3P ^46^ is based on gaseous water, while SPC/E ^47–48^ is based on ice. TIP4P and TIP4P/2005 are 4-point water models, also based on the geometry of TIP3P but with an extra particle to displace the partial charge associated with the oxygen away from the oxygen site. TIP4P, along with TIP3P, is routinely used in biomolecular simulations, whereas the more recently developed TIP4P/2005 model is not yet commonly used. OPC is a recently developed 4-point model with a novel geometry ^45, 49–50^. The latter includes a bond angle of 103.6°, smaller than the gaseous bond angle of 104.5°, and a smaller than expected O-H bond length of 0.8724 Å. The OPC geometry was developed by optimising the point charge arrangement in comparison to quantum mechanical calculations ^45^.

**Figure 4:**
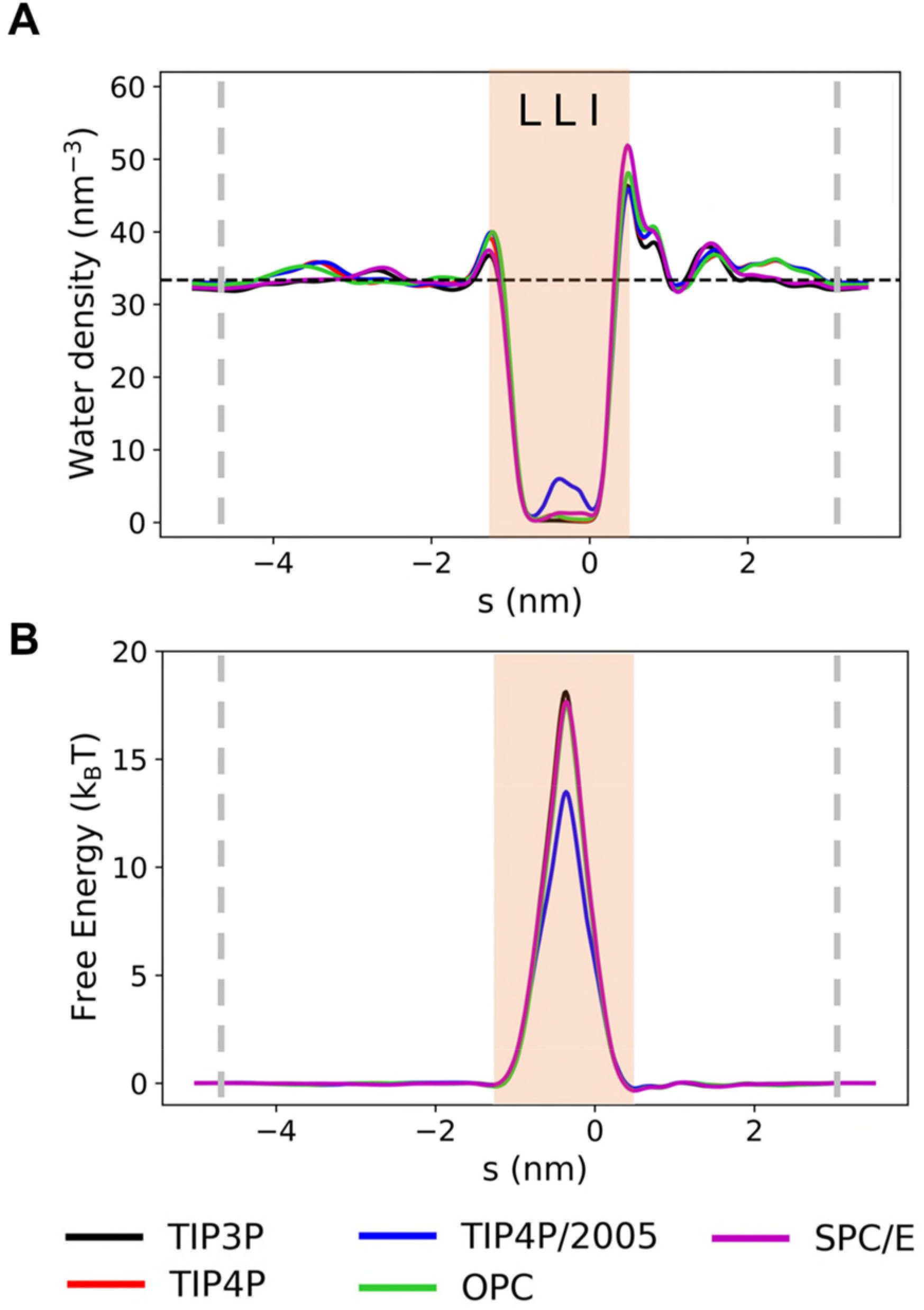
The effect of changing the rigid fixed-charge water model on the water density (**A**) and free energy (**B**) profiles for the wild-type TMEM175 structure. The position of the hydrophobic gate is shaded in pale orange, and the approximate position of the lipid head groups is given by the grey dashed lines. (See also SI Figs. S2, S3 and S4).

For all five additive models, the pore de-wetted as evidenced by the water density profiles (Fig. 4A) and the associated free energy barriers (Fig. 4B). In the case of TIP4P/2005, some water molecules were trapped by the constrictions at the ends of the hydrophobic gate, as discussed above, resulting in a decrease in the free energy barrier. This difference is however just within the limit of the standard deviations (averaged over repeats) for the five water models (see SI Figs. S2, S3 and S4, and therefore we attribute this to stochastic variation rather than a significant difference due to the water model. We therefore conclude that the wetting/de-wetting behaviour is not substantially affected by changes in additive water model, and that these water models are equally valid for investigating the effects of hydrophobic gating. In the absence of direct experimental data, it is difficult to assess whether the water models used are transferrable from bulk to nano-confined (i.e. channel) systems. However, recent experimental studies have attempted to estimate water profiles within the hydrophobic gate of an ion channel ^51^ (but not yet for TMEM175). In the meantime, we can state that the five additive water models explored appear to be equally transferrable when moving from bulk water to water confined in a nanopore.

Comparing the five additive water models in the mutants also produced consistent results. For example, in the case of the AAA (Fig. 5A and SI Fig. S5 & S6) and NNN (SI Fig. S7 & S8) mutants, the hydrophobic gate remains wetted across the five water models. Similarly, in the VVV mutant, the hydrophobic gate de-wets in all cases (Fig. 5B and SI Fig. S9 & S10). Any differences between water model across the free energy profiles are well within the standard deviations and are therefore due to stochastic variation rather than due to the choice of water model (see SI Figs. S6, S8 & S10).

**Figure 5:**
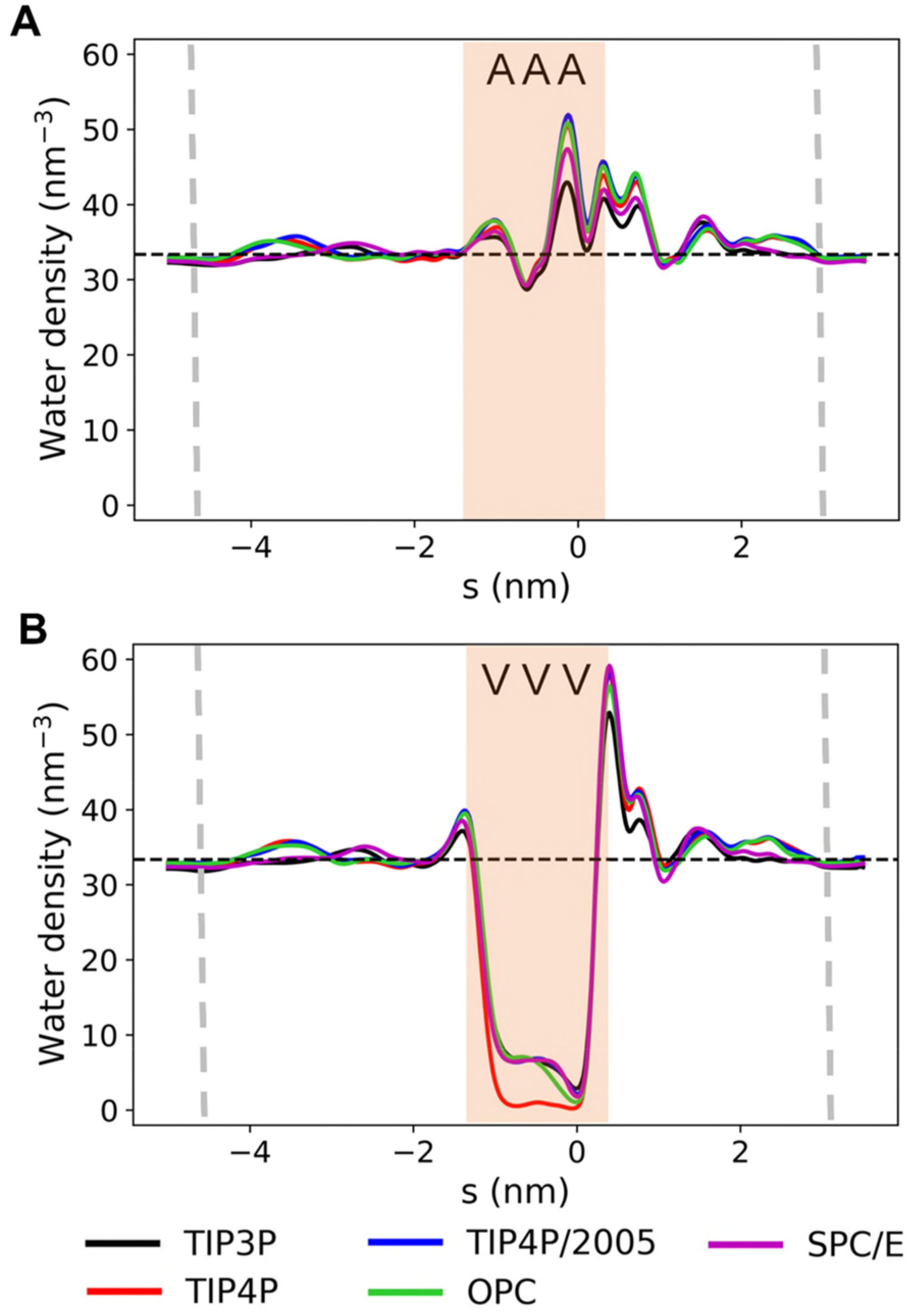
Water density profiles for different rigid fixed-charge water models for the AAA mutant (**A**) and the VVV mutant (**B**). Bulk water density is denoted by the black dashed line. The position of the hydrophobic gate is shaded in pale orange, and the approximate position of the lipid head groups is given by the grey dashed lines. (See also SI Figs. S5–10).

### Effect of a Polarisable Water Model on Wetting/De-wetting

We are especially interested in how polarisable water would behave in the TMEM175 hydrophobic gate structure given the stochastic observation of a ‘trapped’ water (or waters) in this extreme example of nanoconfinement, as seen in our simulations using e.g. the TIP4P/2005 water model (see above and SI Fig. S1B). Experimentally, water in hydrophobic nanoslits exhibits an anomalously low dielectric constant largely reflecting their rotational immobilisation ^39^. Simulation studies have been used to compare a rigid fixed-charge (SPC/E) water model and a flexible polarisable (ABEEM-7P) model for water molecules confined in graphene nanocapillaries with a width ranging from 0.59 to 1.1 nm. In the latter case the polarisable model resulted in more ordered water than with SPC/E ^52^.

Our simulations of the extreme nanoconfinement within the TMEM175 gate (the radius in the middle of the hydrophobic gate is ~0.16 nm for the wild-type (WT) channel, comparable to that of a single water molecule) reveal a difference in the de-wetting behaviour when comparing the polarisable AMOEBA14 water model ^38^ to the five additive water models. This is most clear cut for the WT channel, for which the pore radius profiles within the hydrophobic gate are identical for the five non-polarisable and for the polarisable (AMOEBA) models (SI Fig. S2A). For this channel, the number density of water in the nanocavity in the centre of the hydrophobic gate (Fig. 6A) was ~0 nm^−3^ for TIP4P, ~6 nm^−3^ for TIP4P/2005 and ~15 nm^−3^ for AMOEBA, corresponding to free energies along *s* of ~18, ~13 and ~7 kBT respectively (Fig. 6B). Thus, water within this extreme nanoconfinement (Fig. 7A; the volume of the nanocavity is ~0.04 nm^3^, i.e. just above the volume occupied by a single water molecule in the bulk state, 0.03 nm^3^) is stabilized by 6 to 11 k_B_T (i.e. 15 to 28 kJ/mol) by the use of a polarisable model. Of course, this is an approximate estimate, assuming six repeat simulations provide adequate sampling. For example, FEP calculations could be used to estimate this more exactly.

**Figure 6:**
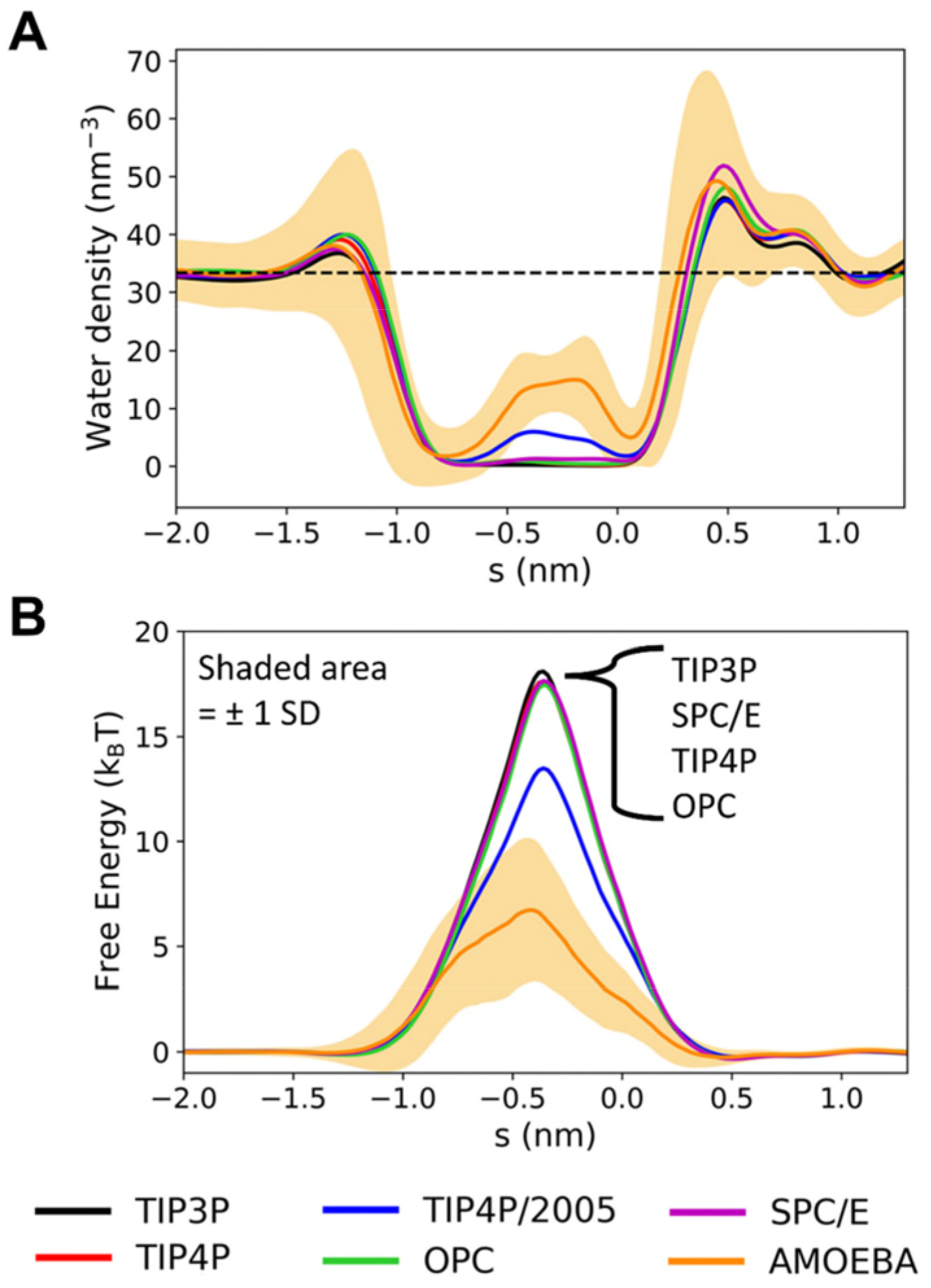
Comparison of water density (**A**) and free energy (**B**) profiles for the TIP4P/2005 fixed-charge and the AMOEBA polarisable water models for the wild-type TMEM175 structure. The shaded regions denote ±1 standard deviation calculated over all simulation frames included in the analysis (discarding the first 10 ns as equilibration), and averaged over all repeats. The dashed black line denotes the value of bulk water density.

**Figure 7:**
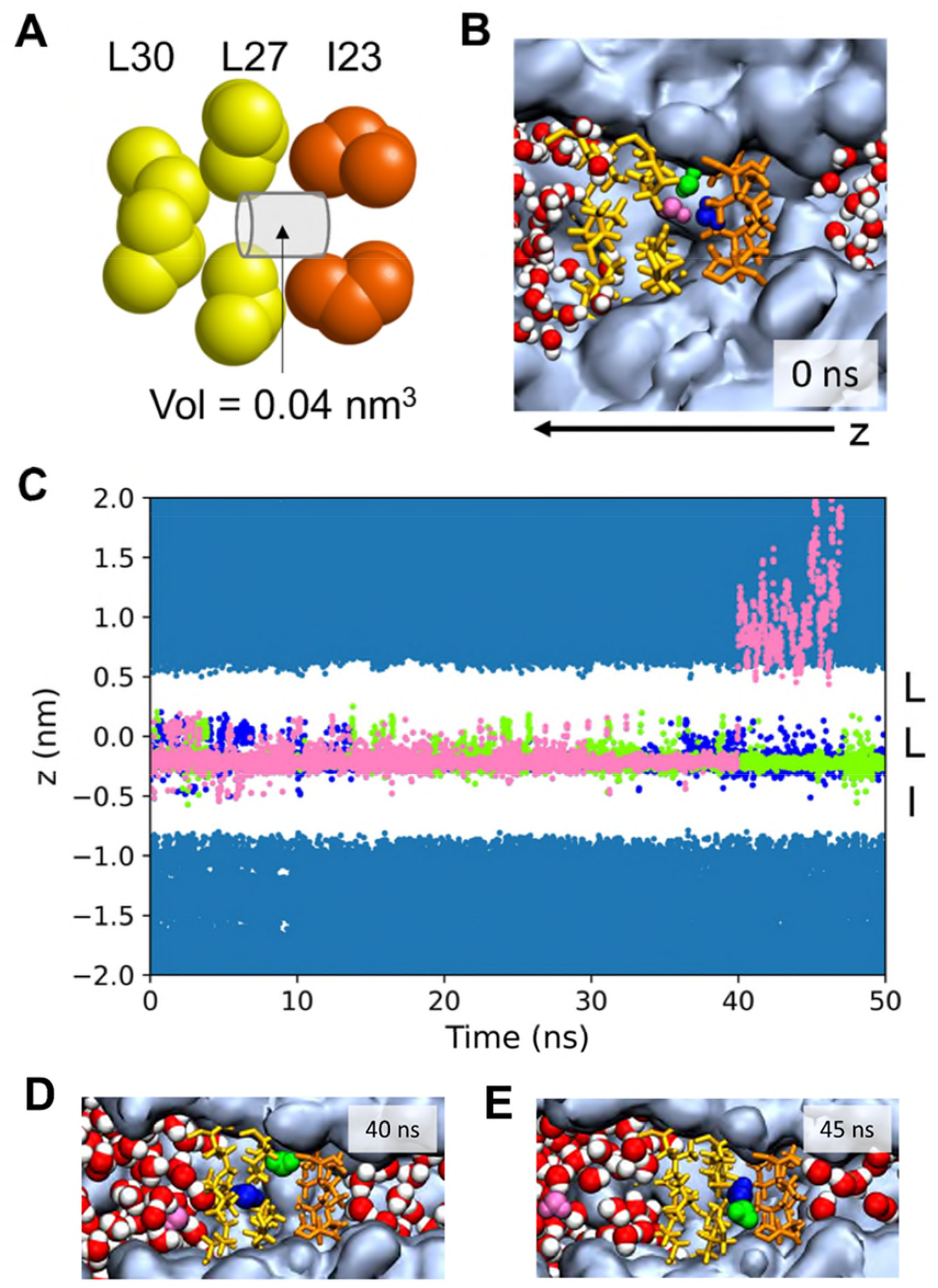
**A** Schematic of the ‘extreme nanocavity’ in the TMEM175 hydrophobic gate, showing the cylindrical approximation to the WT cavity volume. **B** Snapshot of three ‘trapped’ water molecules (coloured blue, pink and green) within the nanocavity which partially hydrate the hydrophobic gate in the AMOEBA simulations. **C** Trajectories of the three trapped waters (coloured as in **B**) for an AMOEBA simulation. The pink water molecule can be seen to leave after ~40 ns. **D and E** Snapshots of the simulation in **C** immediately after and ~5 ns after the pink water molecule has departed from the nanocavity, showing the relaxation of the two remaining water molecules so that they sit between the I32 and L27 sidechain rings. (See also SI Fig. S11).

For the VVV simulations (SI Fig. S9 and S10) the situation was less clear. With this (modelled) mutant pore there was a degree of dependence of the pore radius profile on the water model employed which complicated such detailed comparison.

Overall, this suggests that the use of a polarisable water model has a significant effect on the de-wetting/wetting behaviour of a ‘tight’ hydrophobic gate, and that future simulations of hydrophobic gating in general would benefit from a systematic investigation into the effects of including molecular polarisability.

We can make partial comparisons with other studies. Previous simulation studies of the effect of including polarisability focussed on a less extreme case of nanoconfinement ^27^ involving the gating region of a M25 nanopore derived from the (partially) open state of the 5HT3R channel (PDB 6DG8). For this channel system the time-averaged water density was respectively ~22 nm^−3^ (for TIP3P water), ~16 nm^−3^ (TIP4P/2005) and ~35 nm^−3^ (AMOEBA). This corresponds to an approximate estimate via the Boltzmann equation (where *ρp* and *ρ_NP_* are the water densities for the polarisable and non-polarisable models respectively):

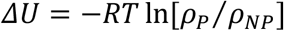

of stabilisation of the nanoconfined water within the hydrophobic gate of ~−1.6 kJ/mol by including polarisation. A comparable estimate for TMEM175 yields a stabilisation of ~−4.2 kJ/mol by including polarisation. An approximate calculation of the relative potential energies of a water dipole in a single molecule sized cavity in dielectrics of 1 (i.e. *in vacuo*) vs. 4 (for a hydrophobic cavity in a protein) yields about −7 kJ/mol stabilisation of the latter case ^53^. Thus, the same trend is seen for the two nanopore systems but the stabilisation by including polarisation is more marked in the extreme nanoconfinement case of the closed hydrophobic gate of TMEM175, which can accommodate ~1-4 water molecules compared with the (partially) open hydrophobic gate in the 5HT3R 6DG8 structure which can more readily accommodate multiple water molecules.

Comparisons with experimental data remain difficult but we note that measurements of the refractive index of nanoconfined water (corresponding to its dielectric constant at optical frequencies ^40^) suggest a decrease in molecular polarisability corresponding to a reduction in refractive index from *n* = 1.33 (bulk) to *n* = ~1.25 for water confined in ~10 nm gaps of hydrophobic interfaces. It would therefore be of great interest to know how the refractive index of water behaves on nanoconfinement scales of < 1 nm.

### Trajectories of De-wetting

We examined the de-wetting behaviour of trapped waters in the hydrophobic nanocavity using the polarisable AMOEBA14 model in a little more detail (Fig. 7 and SI Fig. S11). We tracked the trajectories projected onto the *z* axis of all water molecules initially in the hydrophobic gate region. In two of the three simulations, 3-4 waters were initially present but 1-2 of these exited within the first ~40 ns. In the third repeat the hydrophobic gate was fully de-wetted at the start and remained so for the duration (50 ns) of the simulation (SI Fig. S11).

We also examined the first repeat of these simulations in more detail (Fig. 7). At the start of the simulation three water molecules were present within the central hydrophobic nanocavity. This is divided into two ‘sub-cavities’ one formed by the I and L sidechain rings and one by the L and L sidechain rings. The water molecules switch back and forth between these two sub-cavities on an approximately nanosecond timescale. Just before 40 ns, one water molecule exits from the hydrophobic gate. The remaining two water molecules then continue to switch between the two sub-cavities: sometimes one in each (e.g. Fig. 7D) sometimes both water molecules in one sub-cavity (e.g. Fig. 7E). In the latter case the two waters form a hydrogen bond. Similar behaviour was seen in the second repeat simulation (SI Fig. S11B), during which two water molecules leave early on (< 10 ns) and then two remaining waters switch back and forth dynamically between the same or different nanocavities.

By contrast, in the TIP4P simulations (SI Fig. S1A), in 5 (out of 6) simulations trapped waters were present but were expelled in <15 ns in each case. In the TIP4P/2005 simulations (SI Fig. S1B), in 4 (out of 6) simulations initial trapped water molecules were expelled, whereas in one repeat they mostly remained trapped for the entire simulation. Taken together, these results are consistent with the overall instability of the wetted state of the hydrophobic nanocavity. A metastable state with one or more trapped waters may exist, the exact stability of which is sensitive to the water model employed.

## Conclusions

From the results described above, it can be seen that the broad picture of hydrophobic gating (explored for a closed state of a hydrophobic gate in an ion channel protein) is robust to changes in additive (rigid fixed-charge) water models. However, quantitative details alter when electronic polarisability is included in the simulations. This in turn suggests the need to include consideration of polarisability (see e.g. ^54^ for a recent example) when designing hydrophobic gates or comparable structures into novel nanopores ^14, 55^. Relatively small changes in detailed nanoscale structure (and hence in the local dielectric/polarisability properties) could be used to ‘fine tune’ a hydrophobic gate to e.g. electrowetting.

In this study we have used an X-ray structure of a prokaryotic TMEM175 structure (CmTMEM175 at 3.3 Å resolution; PBD 5VRE ^33^) as a model for investigating the effect of mutating the pore lining and water model on wetting/de-wetting behaviour. CryoEM studies of the human TMEM175 channel ^34^ reveal two possible open and closed structures. Further TMEM175 structures (MtTMEM175) have been solved by X-ray diffraction to 2.4 Å resolution (PDB 6HD8^35^). These differ from the 5VRE CmTMEM175 structure in some respects. In particular, the pore-lining isoleucine (Ile23) in CmTMEM175 is replaced by a leucine (Leu35) in MtTMEM175. Together these structures indicate conservation of a hydrophobic nanoscale constriction in the transmembrane pore.

The current studies could be extended in the future by applying polarisable (AMOEBA) simulations to characterize water behaviour in a range of channel structures selected to have hydrophobic gates of differing dimensions ^19^. This could enable us to explore the effect of the *degree* of nanoconfinement on the extent of anomalous water behaviour. Combining recent computational (this study) and experimental (e.g. ^51^) approaches would therefore allow us to exploit biological nanopores/channels to characterise the effects of confinement on water behaviour on smaller scales than those currently addressed (down to ~1.5 nm) by studies of nanofabricated devices ^39^. In the limiting case, this may enable us to use biological systems to examine the behaviour of water under extreme nanoconfinement with just one or two water molecules present. This is of relevance to a number of areas of enquiry, especially given the continued interest in the behaviour of water confined in nanoporous environments ^56–57^.

## Methods

### Protein structure, preparation

The prokaryotic TMEM175 structure from *Chamaesiphon minutus* (CmTMEM175 PBD: 5VRE) was run through the WHATIF server (https://swift.cmbi.umcn.nl/whatif/) to fix any missing sidechains ^58^. The three rings of pore-lining residues forming the hydrophobic gate in the wild-type (Ile23, Leu 27 and Leu30 on each of the four subunits) were then mutated in PyMol (http://www.pymol.org) to three rings of alanines, asparagines or valines to form the three mutant structures (‘AAA’, ‘NNN’ and ‘VVV’ structures).

### Non-polarisable simulations

TMEM175 structures were embedded in a POPC (1-palmitoyl-2-oleoyl-*sn*-glycero-3-phosphocholine) bilayer using the MemProtMD and CG2AT protocols ^59–60^. This uses a multi-scale methodology to solvate the system with ~0.15 M NaCl, self-assemble the lipid bilayer and equilibrate the system. Production runs involving the additive water models were performed using GROMACS ^61^ version 2016.3 with the OPLS all-atom protein force field with united-atom lipids ^62^. Each simulation was run for 100 ns with an integration time step of 2 fs. For the CHAP analysis (see below), the first 10 ns were discarded as equilibration. 6 repeats were run for each protein structure and water model combination. The temperature was maintained at 310 K using the V-rescale thermostat ^63^ and a time coupling constant of 0.1 ps. The pressure was maintained at 1 bar using the semi-isotropic Parrinello-Rahman barostat ^64^ and a time coupling constant of 1 ps. The Verlet cut-off scheme was used, along with the Particle Mesh Ewald method for electrostatics ^65^. The LINCS algorithm was used to constrain bonds ^66^.

Notably, the positions of the protein backbone atoms were position restrained with a force constant of 1000 kJ mol^−1^ nm^−2^, however protein side chains were allowed to relax. We chose to apply these position restraints because we did not want our overall simulation protein structure to deviate from the original experimental coordinates. It was important to do this because we specifically wanted to assess how changes in the protein pore lining (i.e. via mutants) and water model affected the wetting/de-wetting behaviour within a given overall protein geometry.

### Polarisable simulations

These were performed using a similar method to that described in Klesse et al. ^27^ (https://github.com/Inniag/openmm-scripts-amoeba). The wild-type and VVV mutant structures were embedded in a DOPC bilayer and solvated with ~0.15 M NaCl solution using the bilayer self-assembly method outlined for the non-polarisable simulations. A 10 ns equilibration simulation was then run using GROMACS with the non-polarisable CHARMM36m force field. Polarisable production runs were performed using OpenMM ^28^, and lasted for 50 ns for the wild-type structure and 35-50 ns for the VVV mutant structure. The AMOEBA force field was used for all atoms in the simulation i.e. for the protein ^67^, ions ^68^ and lipids ^69^, together with the AMOEBA14 water model ^38^. The r-RESPA algorithm was used with an outer timestep of 2 fs and an inner timestep of 0.25 fs. The pressure was maintained at 1 bar using the Monte Carlo barostat and the temperature was maintained at 310 K using the Andersen thermostat. The Cα backbone atoms were position restrained with a force constant of 1000 kJ mol^−1^ nm^−2^ for both the equilibration and production runs. There were 3 repeats for each structure.

### Analysis

Profiles for pore radius, water density and free energy were obtained using ChAP (www.channotation.org)^24^. A bandwidth of 0.14 nm was used for calculating the water density through the pore. The free energy profiles were obtained via:

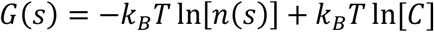

where *G*(*s*) is the free energy of wetting/de-wetting, *n*(*s*) is the water density, s is the coordinate passing down the central axis of the pore (approximately perpendicular to the bilayer), *T* is the temperature, *k_B_* is the Boltzmann constant and *C* is an arbitrary constant set so that *G*(*s*) = 0 outside the pore. Having first discarded the first 10 ns of the simulation as equilibration, these profiles were averaged over all remaining frames of the simulation, and then averaged over all repeats as appropriate.

The positions of individual water molecules were calculated with the aid of MDAnalysis ^70–71^. It should be noted that the protein is oriented such that z axis is approximately antiparallel to the local pore *s* defined by CHAP and so for all practical purposes *z* = -*s*. Figures were created using VMD ^72^ and Matplotlib ^73^ (<u>10.5281/zenodo.592536</u>).

## Author Information

Corresponding authors:

## Acknowledgement

This work was supported by grants from EPSRC (EP/R004722/1; EP/V010948/1; EP/R029407/1), BBSRC (<u>BB/N000145/1</u>) and Wellcome (208361/Z/17/Z). The authors thank Miguel A. Gonzalez for discussions on water model implementation.

---

**Figure S1:**
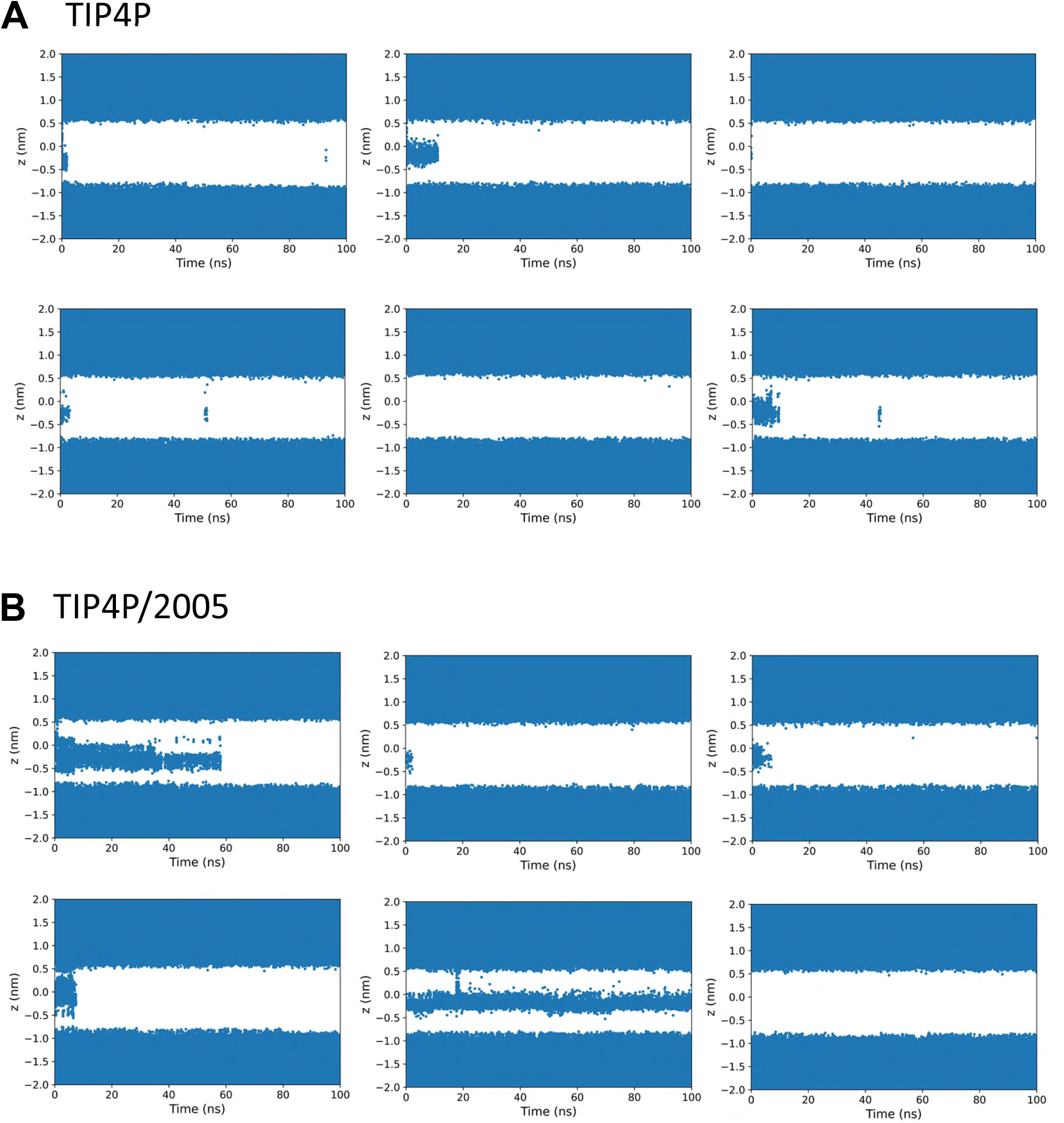
The trajectories of water molecules in and around the hydrophobic gating region (centred at *z* = 0) shown projected onto the *z* axis for each of the 6 repeats of the **A** TIP4P and **B** TIP4P/2005 simulations of the wild-type channel.

**Figure S2:**
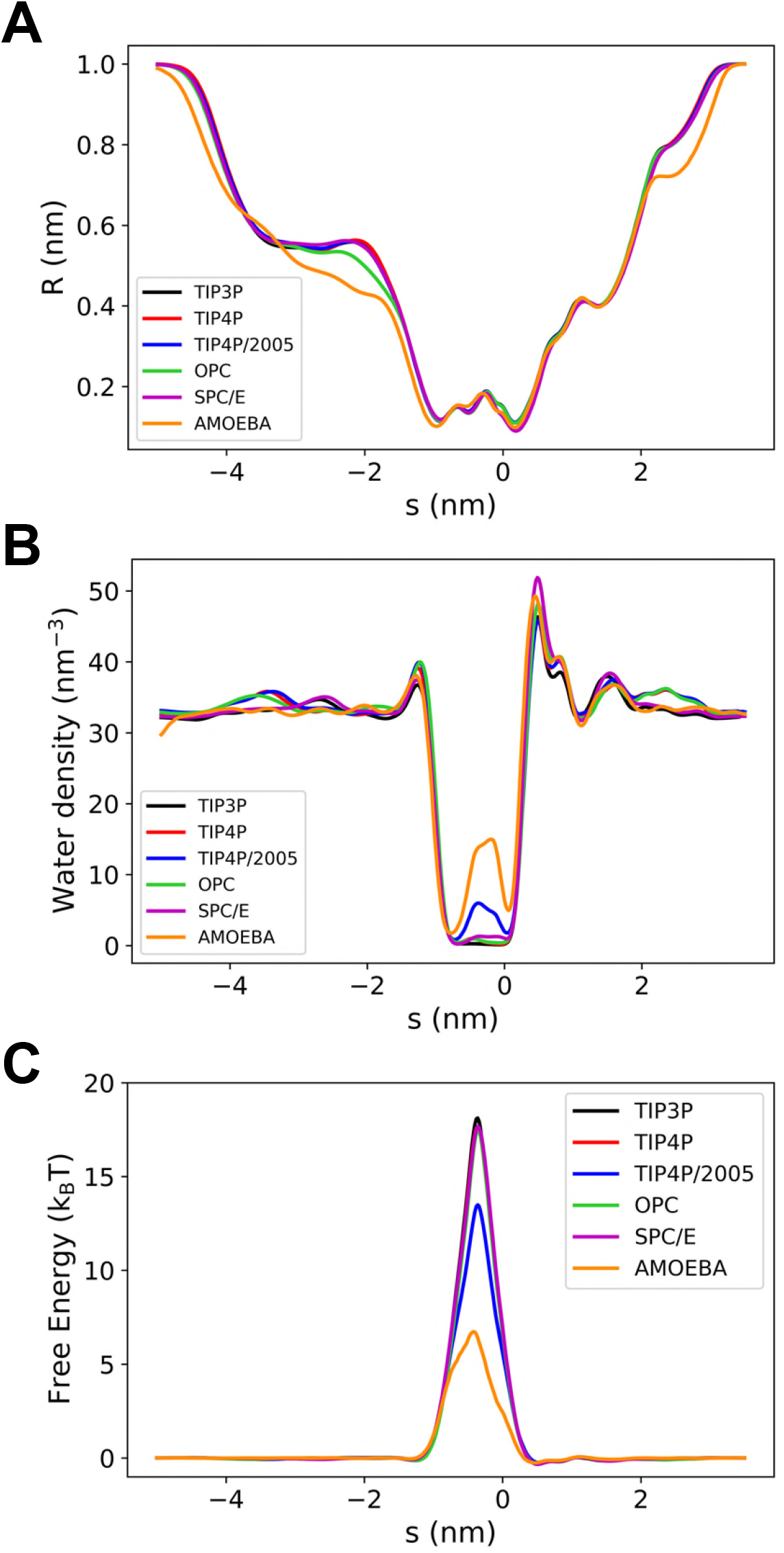
Profiles for the wild-type channel as a function of water model: **A** radius, **B** water density, **C** water free energy.

**Figure S3:**
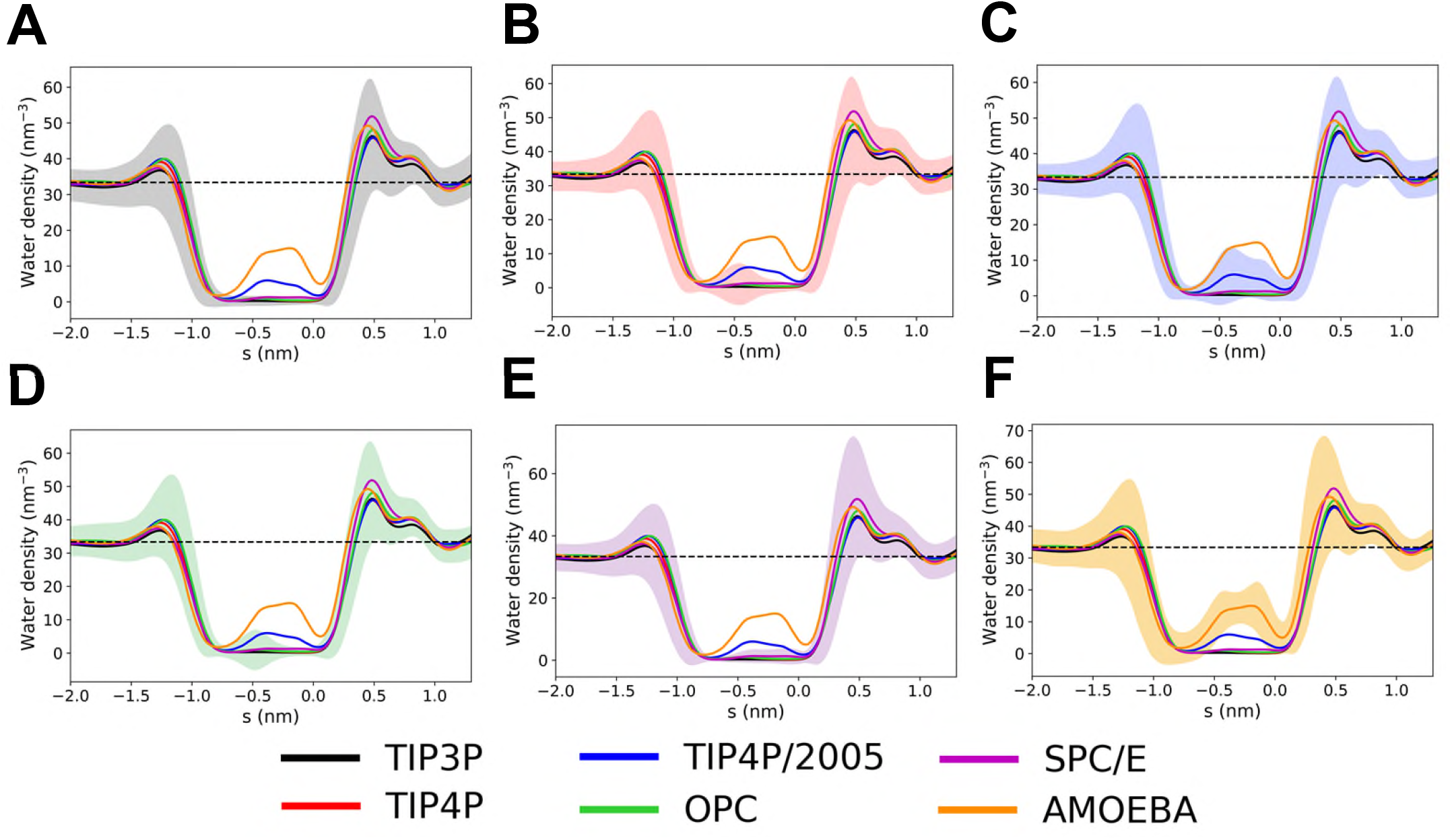
Water density profiles for the wild-type channel, showing the standard deviation for each water model (shaded regions): **A** TIP3P, **B** TIP4P, **C** TIP4P/2005, **D** OPC, **E** SPC/E and **F** AMOEBA.

**Figure S4:**
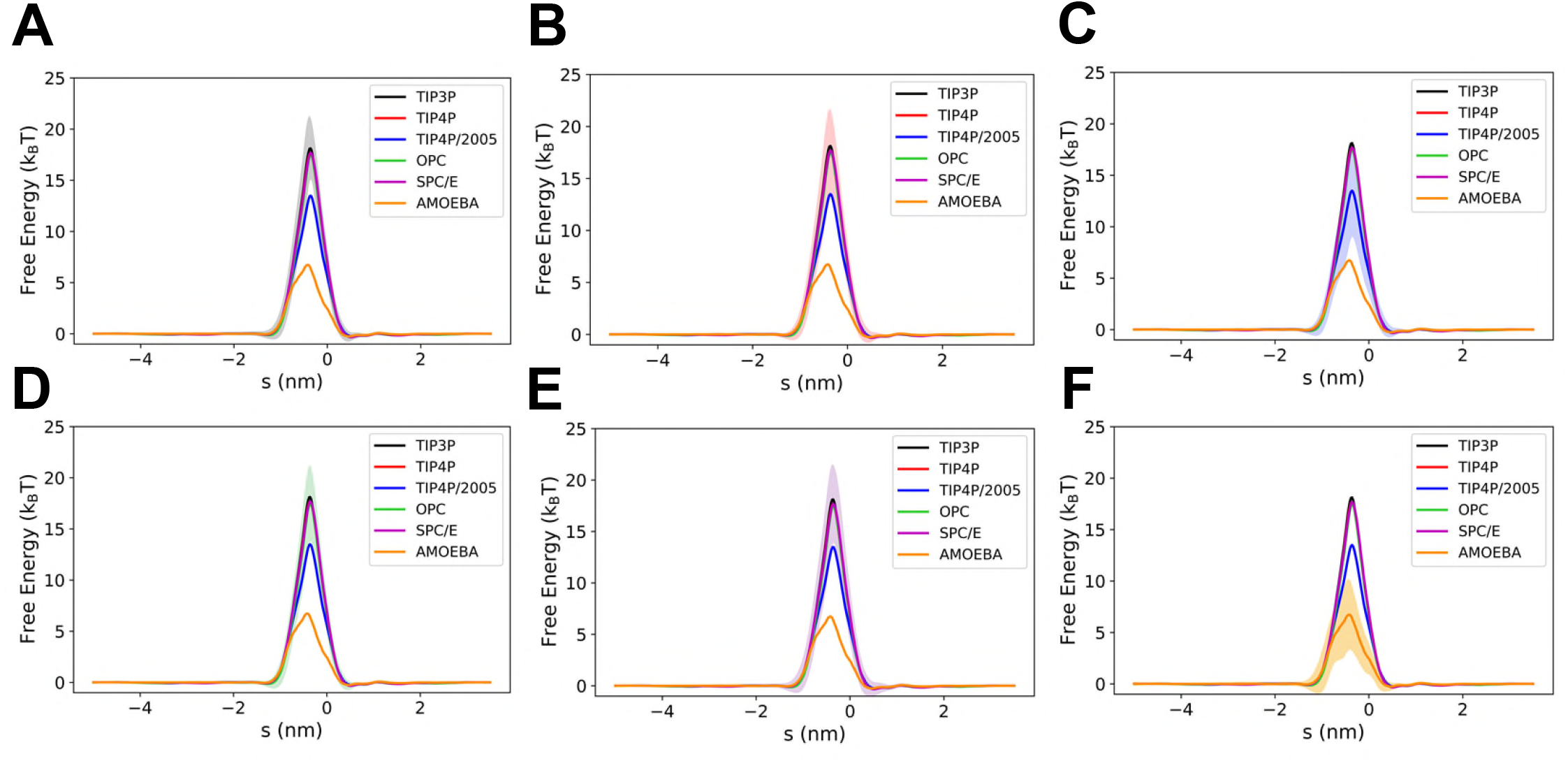
Water free energy profiles for the wild-type channel, showing the standard deviation for each water model (shaded regions): **A** TIP3P, **B** TIP4P, **C** TIP4P/2005, **D** OPC, **E** SPC/E and **F** AMOEBA.

**Figure S5:**
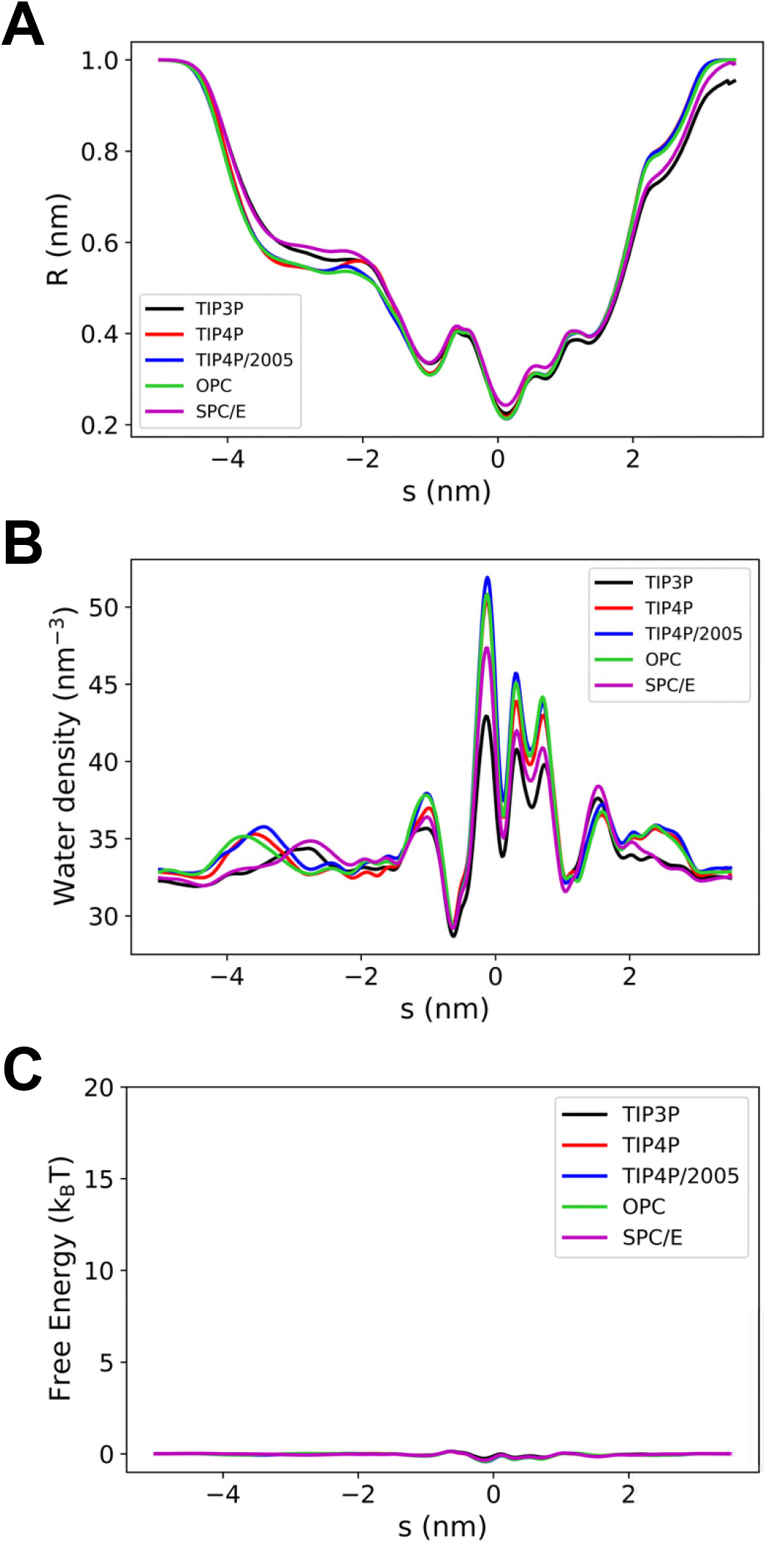
Profiles for the AAA mutant channel as a function of water model: **A** radius, **B** water density, **C** water free energy.

**Figure S6:**
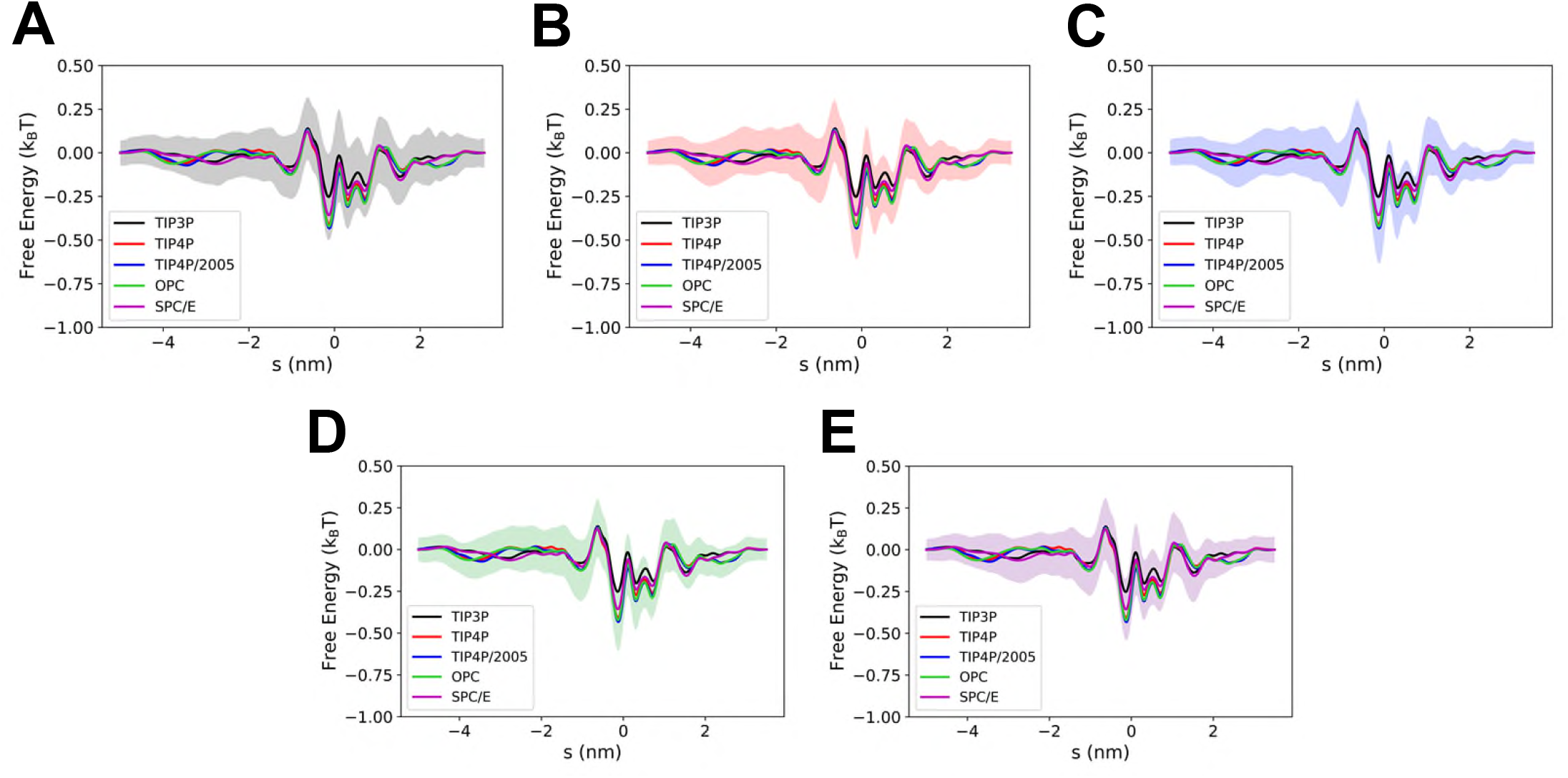
Water free energy profiles for the AAA mutant channel, showing the standard deviation for each water model (shaded regions): **A** TIP3P, **B** TIP4P, **C** TIP4P/2005, **D** OPC, and **E** SPC/E.

**Figure S7:**
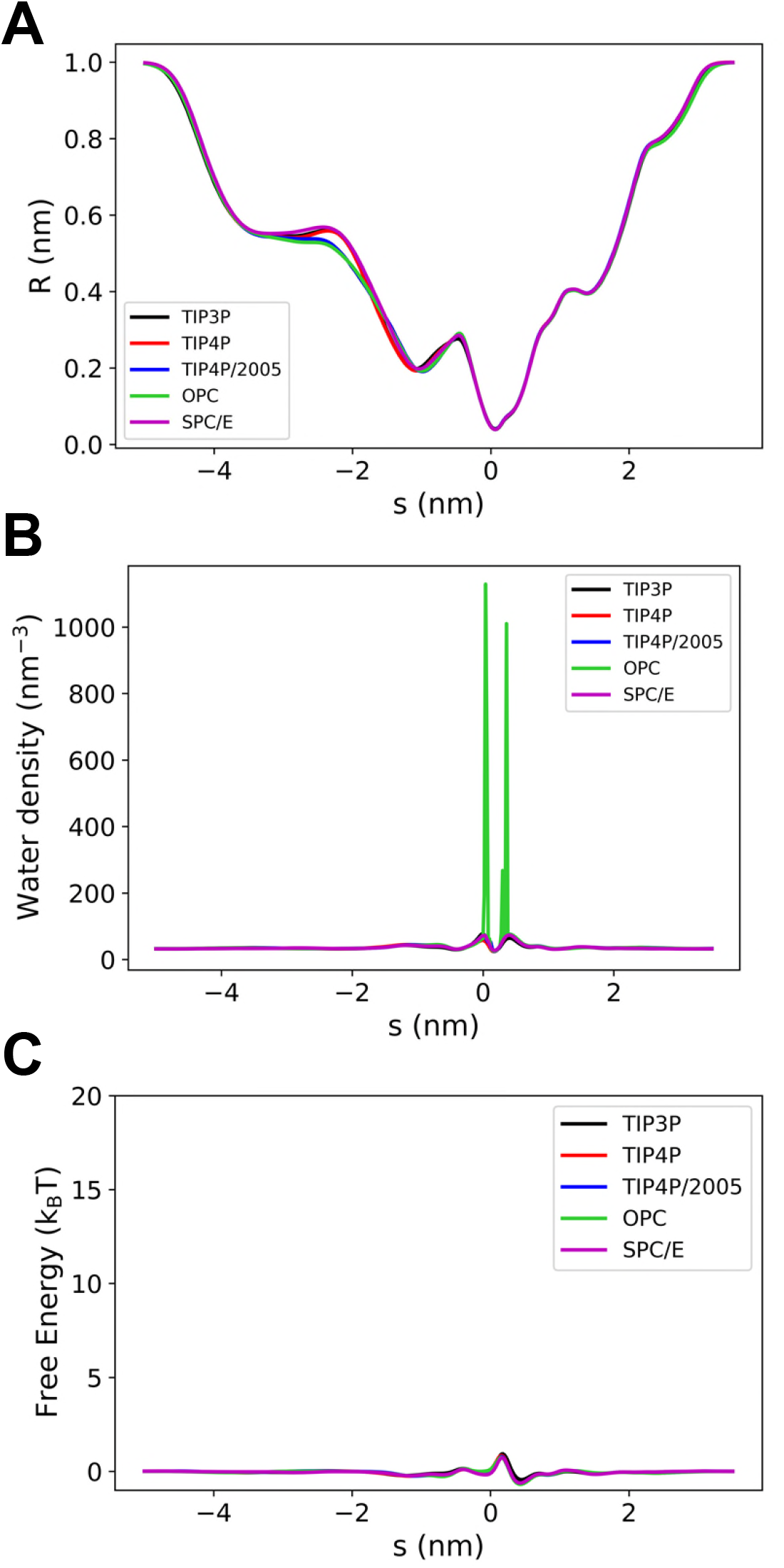
Profiles for the NNN mutant channel as a function of water model: **A** radius, **B** water density, **C** water free energy.

**Figure S8:**
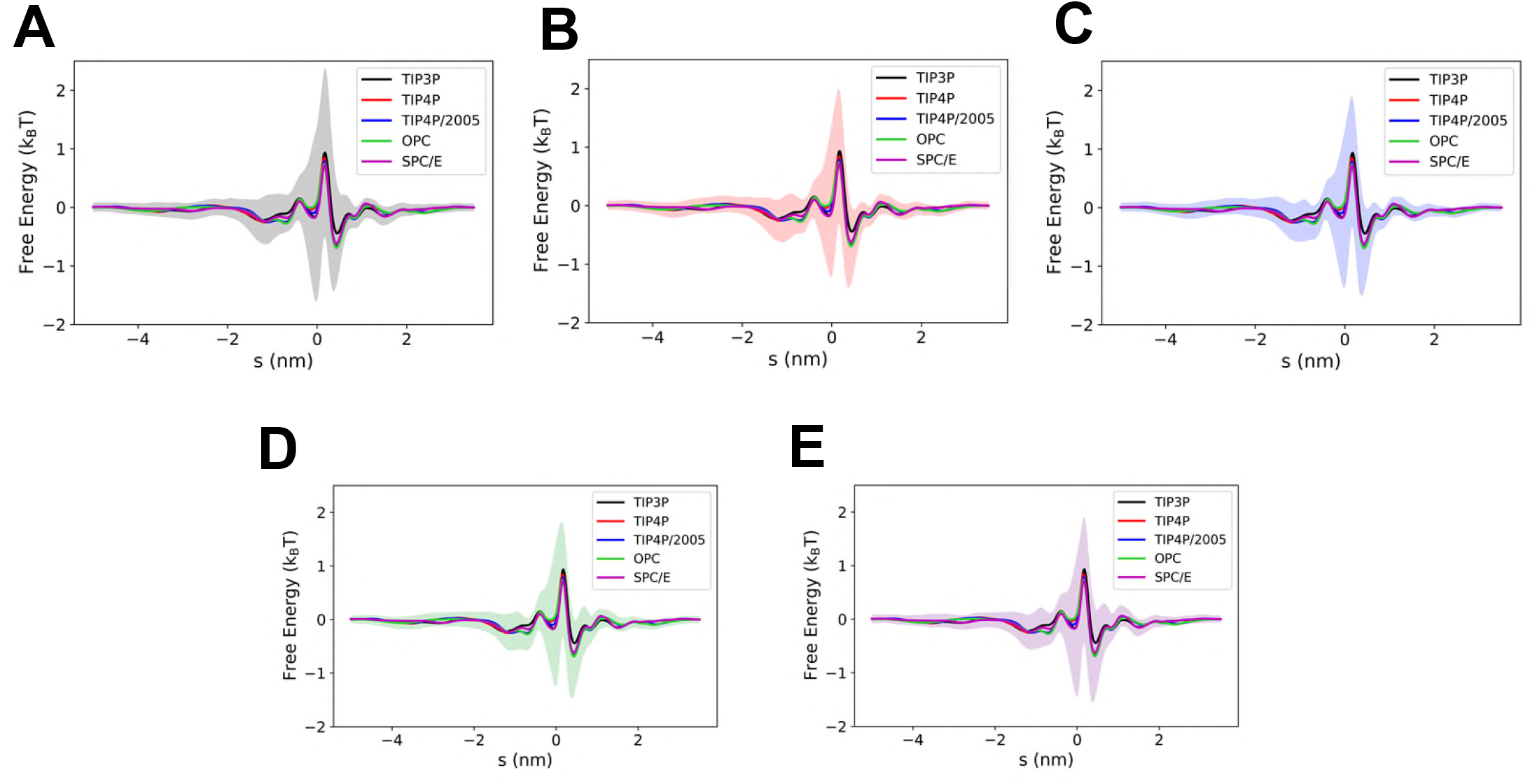
Water free energy profiles for the NNN mutant channel, showing the standard deviation for each water model (shaded regions): **A** TIP3P, **B** TIP4P, **C** TIP4P/2005, **D** OPC, and **E** SPC/E.

**Figure S9:**
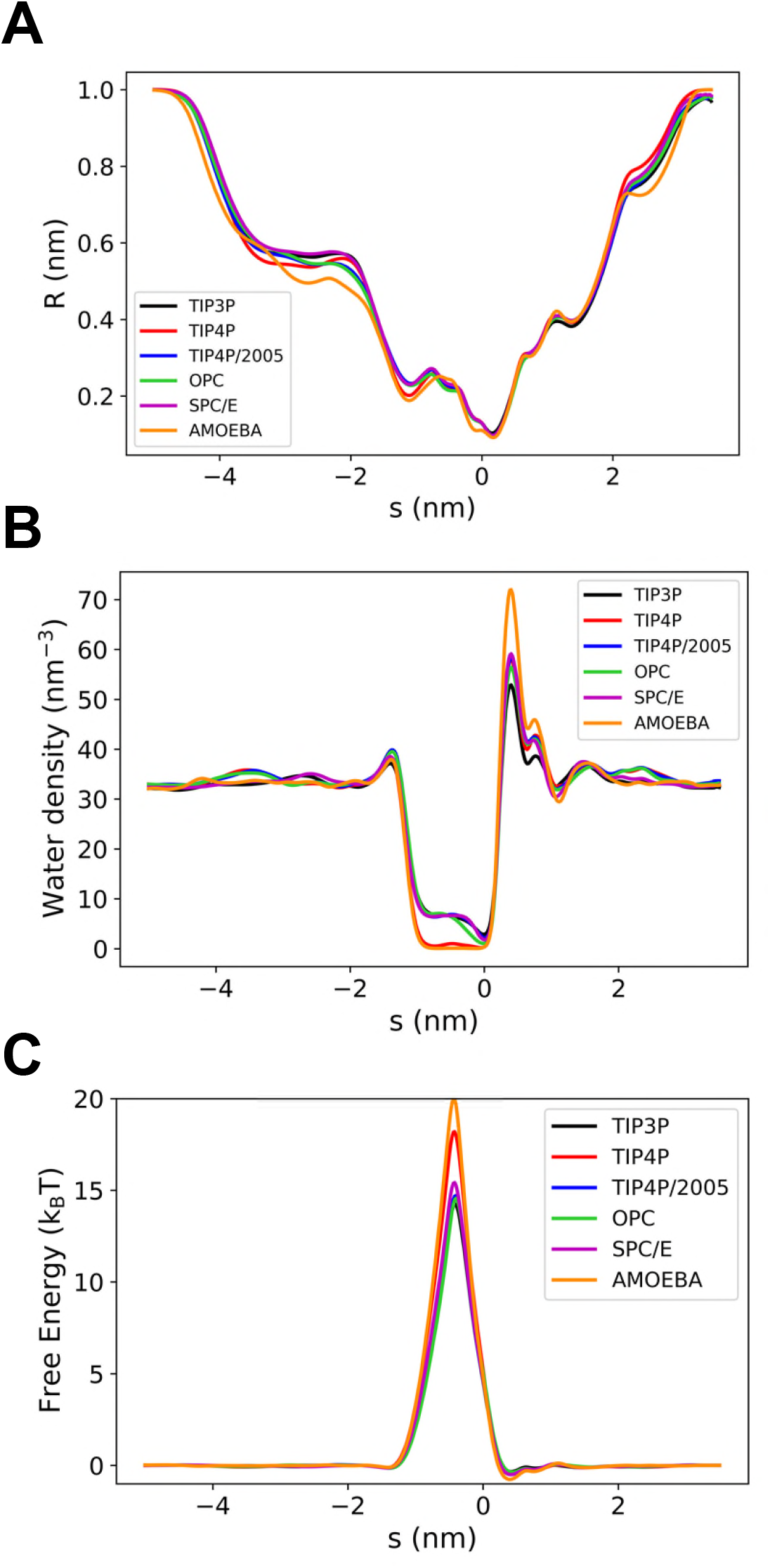
Profiles for the VVV mutant channel as a function of water model: **A** radius, **B** water density, **C** water free energy.

**Figure S10:**
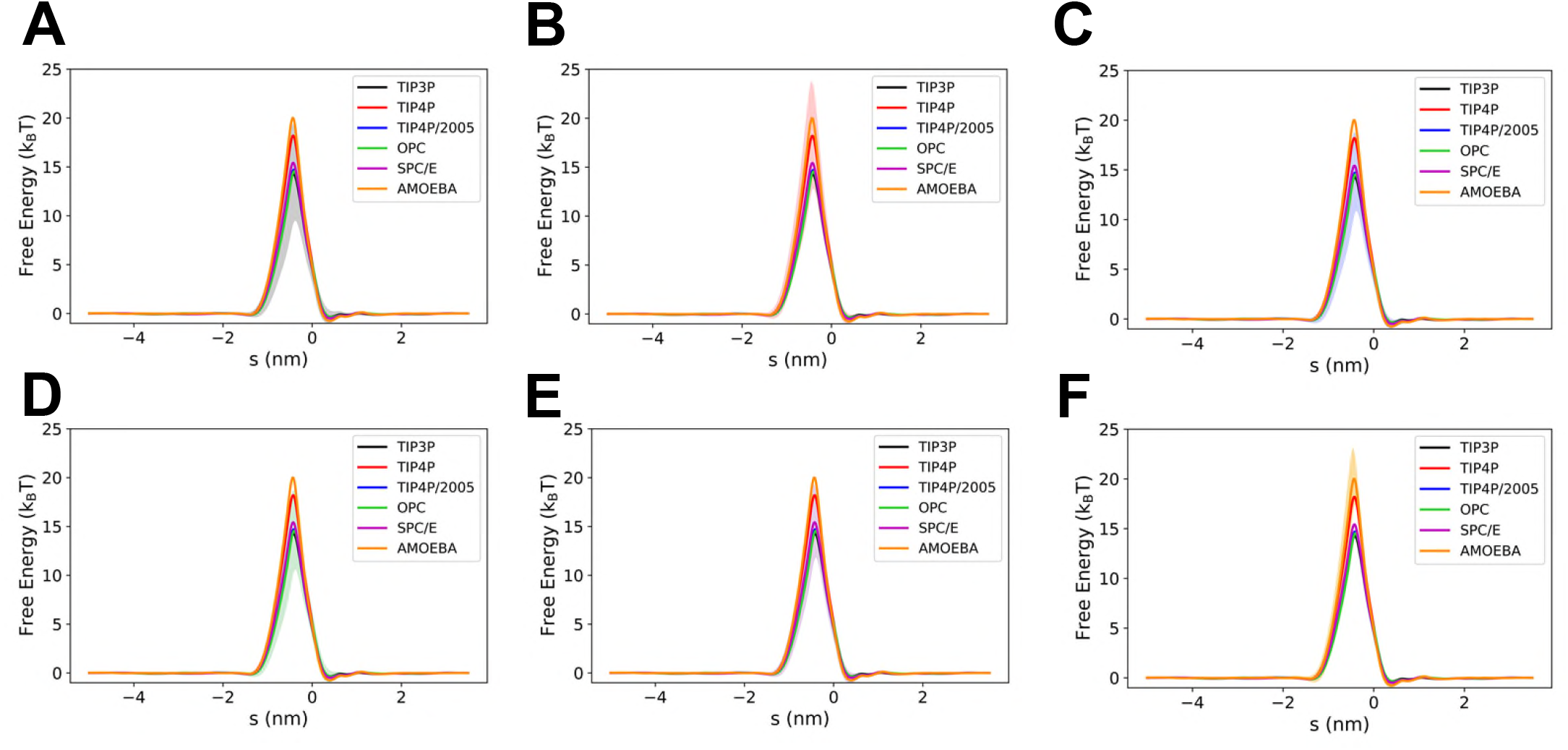
Water free energy profiles for the VVV mutant channel, showing the standard deviation for each water model (shaded regions): **A** TIP3P, **B** TIP4P, **C** TIP4P/2005, **D** OPC, **E** SPC/E, and **F** AMOEBA.

**Figure S11:**
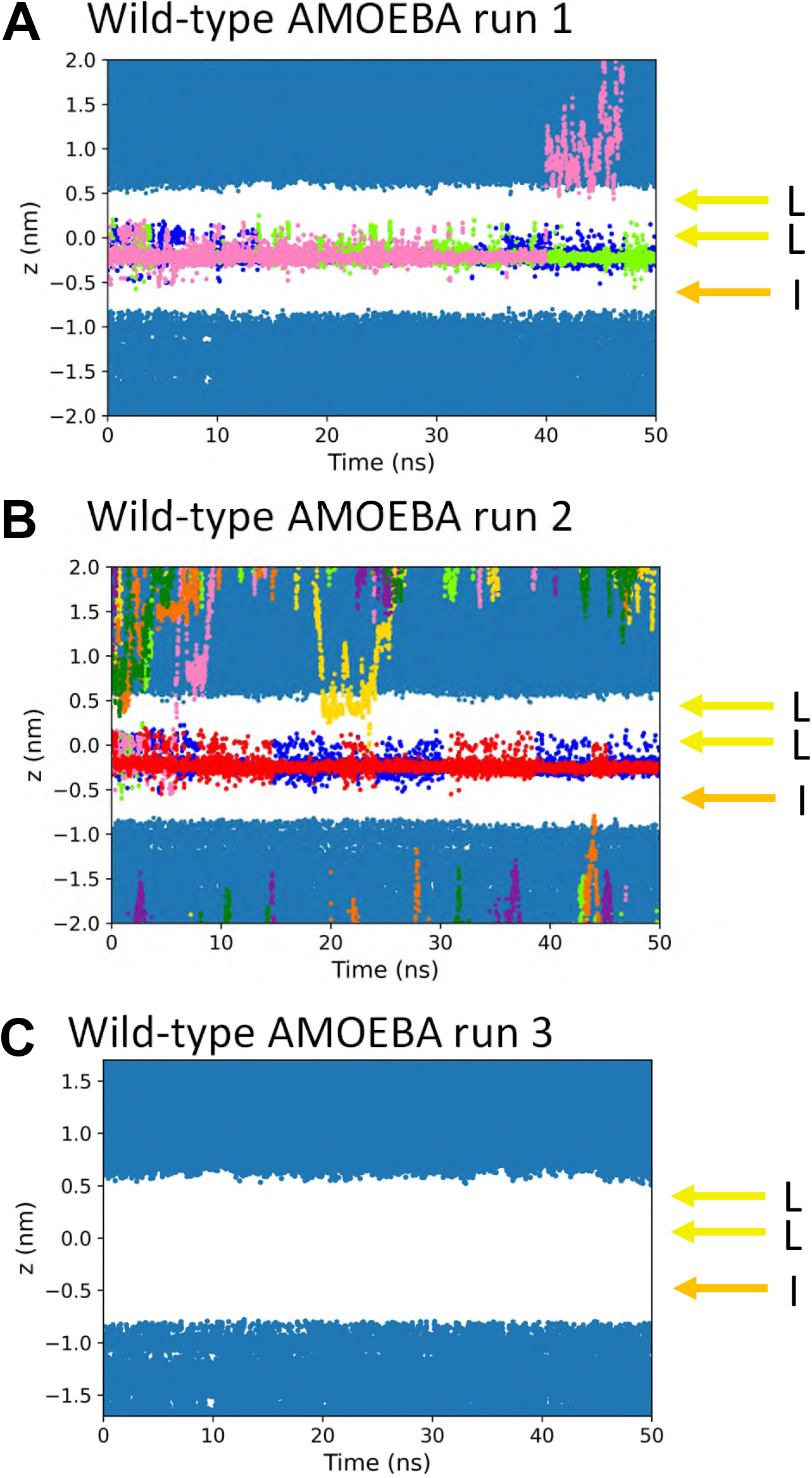
The trajectories of water molecules initially inside the gating region (bright blue, green, red, pink) are shown projected onto the *z* axis (note that *z* is approximately equivalent to –*s*, the pore axis used in the previous figures) within the region of the hydrophobic gate for each of the 3 repeats of the AMOEBA simulation of the wild-type channel. Water molecules which are situated outside the gating region are shown in dark blue, and those which approach the gating region during the simulation are depicted in other colours. The yellow and orange horizontal arrows show the z-coordinates of the centroid of the hydrophobic L and I rings which form the hydrophobic nanocavity.

